# Transcriptional signals of dedifferentiation in human cancer

**DOI:** 10.1101/2022.11.28.518191

**Authors:** Gerda Kildisiute, Maria Kalyva, Rasa Elmentaite, Stijn van Dongen, Christine Thevanesan, Alice Piapi, Kirsty Ambridge, Elena Prigmore, Muzlifah Haniffa, Sarah A. Teichmann, Karin Straathof, Isidro Cortés-Ciriano, Sam Behjati, Matthew D. Young

## Abstract

As normal cells transform into cancers, their cell state changes (or “dedifferentiates”), which may drive cancer cells into a stem-like or more primordial, foetal or embryonic cell state. Here, we used single cell atlases to study dedifferentiation in transcriptional terms across a wide spectrum of adult and childhood cancers. At the level of the whole transcriptome, we find that adult cancers rarely return to an embryonic state, but rather that a foetal state is a near-universal feature of childhood cancers. We extend these bulk transcriptomic findings to a single cell resolution analysis of colorectal and liver cancers, confirming the lack of reversion to a primordial state in adult tumours and the retention of foetal signals in childhood cancers. Our findings provide a nuanced picture of dedifferentiation in these two groups of neoplasms, indicating cancer-specific rather than universal patterns of dedifferentiation pervade adult epithelial cancers.

---

Malignant transformation is underpinned by changes of the cell state towards a less differentiated or stem-like cell state, referred to as dedifferentiation. Consequently, cancers may broadly retain the cell state of origin or assume a dedifferentiated state, of tissue-specific foetal or primordial embryonic cells. Which cell state - primordial or otherwise - a cancer assumes is a fundamental question of cancer biology as it provides a net readout of the consequences of cancer formation.

The cancer cell state can be studied using transcriptional readouts that represent cellular phenotypes and differentiation states. The key challenge in such an analysis is defining cellular states, such as stemness, embryonicness, fetalness, etc., in quantitative molecular terms. One approach is to use appropriate mRNA signals, identified using unsupervised clustering or pattern recognition methods applied to single cell atlases, that embody the state the cancer may (de)differentiate towards. A number of studies have used this approach to investigate “stemness” across the entire spectrum of human cancer^1–3^, for example, by measuring stemness signals derived from *in vitro* differentiating human embryonic stem cells^4^. These studies indicate that human cancer transcriptomes resemble transcriptional modules of *in vitro* differentiating embryonic stem cells. Whether correlation of such modules represents a global transcriptional transformation towards an antenatal state remains unknown. Furthermore, it is conceivable that other developmental states, such as gastrulation, foetal tissues or post-natal stem cells, may also contribute to the differentiation state of cancer transcriptomes.

Recent efforts, referred to as the Human Cell Atlas project^5^, have begun to provide transcriptional definitions of human cells across all stages of development for multiple tissues using single cell mRNA sequencing. These reference transcriptomes provide the opportunity to study dedifferentiation using a more complete and nuanced set of reference states than has been possible in the past.

We approach the quantification of cancer cell state by probing cancer bulk transcriptomes for evidence of single-cell-defined human reference signals using a previously established method^6^. This method, which is conceptually similar to deconvolution, quantifies the extent to which reference signals explain the observed expression profile of a bulk transcriptome. Crucially, this approach also estimates the fraction of the bulk transcriptomic profile unexplained by the provided reference, which together with goodness of fit metrics, assesses the extent to which reference signals account for bulk transcriptomes.

The human reference cell types we included in each analysis encompassed the entire spectrum of human tissue development: human pre-gastrulation epiblast and hypoblast cells^7^; cells representing the three germ layers, endoderm, mesoderm and ectoderm in their earliest stages^8^; and tissue-specific fetal^9^ and adult cells^10^ (**Table S1**). We applied this combined reference to a wide spectrum of childhood and adult solid tissue cancers^11–16^ (**Table S2**). We then extended these results using high resolution atlases of specific tissues together with single cell cancer transcriptomes for two types of adult epithelial cancers.

The starting point of our analysis was to test whether our approach captured similar information to existing measures of “stemness” in human adult cancer. We compared previously published stemness scores that build on gene sets to a broader signal of stemness, the transcriptomes of human pre-gastrulation embryo cells (hypoblast and epiblast). Despite these differences in underlying methodology, we observed that stemness score and our measure, the fraction of each cancer bulk transcriptome explained by human pre-gastrulation embryo cells, correlated strongly (pearson correlation 0.45, **Figure S1**). However, we also found that only a small fraction of each cancer transcriptome could be explained by early embryonic signals (mean goodness of fit = 0.25; mean fraction of unexplained signal = 0.52), which was particularly apparent when compared to positive control transcriptomes (blastoid transcriptomes^17^). Examining which genes drove the stemness score, we found that genes associated with S and G2M phases of the cell cycle were significant drivers of stemness (**Figure S1**). Consistent with this observation, we found stemness scores were higher in dividing than non-dividing tissues (**Figure S1**). Overall, these results suggest that proliferation is a key driver of previously reported stemness scores. This then raises the question whether cancers exhibit tissue-specific signals of dedifferentiation to an antenatal state beyond proliferative signals.

Accordingly, we re-examined bulk transcriptomes by progressively expanding the reference we used for comparison, which enabled us to assess at which point cancer transcriptomes were most completely accounted for. In successive iterations of the analysis, we expanded the reference of pre-gastrulation cells with the following human cell atlases: gastrulation embryo; foetal tissues; and adult tissues. For the control population (blastoids), the vast majority of the bulk transcriptomic signal was explained by the early embryo and gastrulation reference, even when 658,368 foetal and post-natal cells were provided (**Figure 1A-B, S2-4**). By contrast, very little of the early embryonic signals was retained by solid cancers once tissue specific references were available (**Figure 1A-B, S2-4**). Taken together, these results indicate that while cancer cells may share functional features with embryonic stem cells, the cancer transcriptome does not return to an embryonic state.

**Figure 1.**
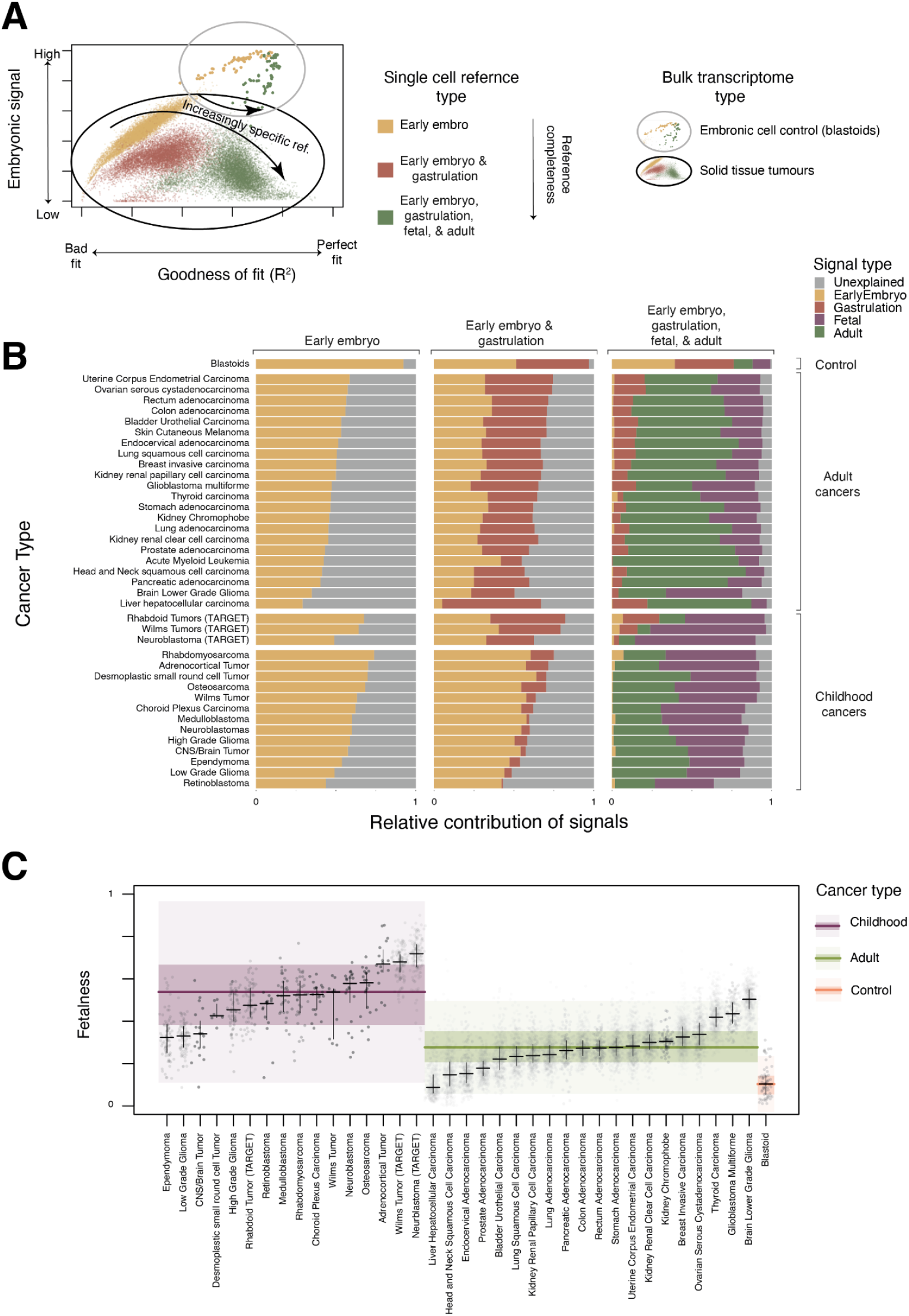
Pan-cancer analysis of differentiation state from pan-tissue single cell reference atlases. **A. Fit quality and embryonic signal of bulk transcriptomes with increasingly complete reference atlases:** Fractional contribution of embryonic reference (y-axis, early embryo + gastrulation for full reference, early embryo otherwise) in explaining bulk transcriptomes (dots) as a function of goodness of fit (x-axis, pseudo R-squared) when fit using single cell reference consisting of cells from the early embryo (yellow), early embryo and gastrulation (dark red), or early embryo, gastrulation, foetal and mature pan-tissue reference (green). Bulk cancer transcriptomes are circled in black and genuinely embryonal controls (blastoids) are circled in grey. **B. Relative contribution of references to explaining the bulk transcriptomes of a range of adult and childhood cancers:** Average relative contribution of early embryo (yellow), gastrulation (dark red), fetal (purple), and adult (green) single cell reference populations in explaining bulk transcriptomes (y-axis) for different combinations of these references (x-axis, labels at top). Bulk transcriptomes are organised by source (labels on right). **C. Childhood cancers have a stronger foetal contribution than adult cancers or control populations:** Relative contribution of foetal reference (y-axis) in explaining bulk cancer transcriptomes (dots), when provided a complete set of early embryo, gastrulation, foetal, and adult single cell references. Bulk transcriptomes are split into childhood (purple), adult (green), and blastoid control (orange) and then by cancer type (x-axis). Distributions are summarised by median (horizontal lines), 1st and 3rd quartiles (horizontal lines for cancer types, shaded coloured areas for childhood/adult/control), and 1.5 times inter-quartile range (light shaded areas).

While embryonic signal seemed to have no discernable biological meaning once more mature references were available, we found a strong and consistent difference between the amount of retained fetal signal, or “fetalness”, in childhood versus adult tumours (**Figure 1C**). This finding extends previous work in renal tumours that described a fetal-like transcriptome in childhood, but not adult kidney tumours^18^. One limitation of our approach is that pan-tissue human atlases, which we used in our analyses, inevitably lack the granularity of annotation and resolution of dedicated, tissue-specific cell atlases. Although our pan-tissue analyses are sufficient to make general statements about stemness, fetalness, etc, they are insufficient to make detailed comparisons to specific cell types. Therefore, in a second part of our analyses we sought to study particular tumours utilising detailed tissue-specific cell atlases and well as single cancer cell transcriptomes for validation of our findings. For this analysis, we focused on liver and colorectal cancer, as both developmental and adult cell atlases of normal liver and colorectal tissues are available.

Hepatocellular carcinoma (HCC) is the most common adult liver cancer. It is an epithelial cancer that arises from hepatocytes, often in the context of chronic liver disease. The precise differentiation state of hepatocellular carcinoma has not been established, compounded by an ongoing debate about the nature of hepatobiliary stem cells. It is noteworthy that some hepatocellular carcinomas resume expression of foetal albumin (alpha-fetoprotein, AFP), which may represent dedifferentiation of some cancers towards a fetal hepatic state^19^. We assessed 360 hepatocellular carcinoma bulk transcriptomes from 3 cohorts, as well as 64 bulk transcriptomes of the childhood liver cancer, hepatoblastoma (HB)^11,12,15^. Using a highly detailed combined reference map of adult and foetal liver^20–22^, together with the pre-gastrulation and early embryo references^7,8^, we asked which reference best explained the bulk transcriptomes state. As with our pan-tissue analysis (**Figure 1**), the reference foetal and adult liver cells accounted for the majority of the transcriptome in hepatocellular carcinoma and hepatoblastoma (**Figure 2A, S5**). The differentiation state of hepatocellular carcinoma bulk transcriptomes was therefore most completely represented by liver cells, but not by more primordial human cell populations.

**Figure 2.**
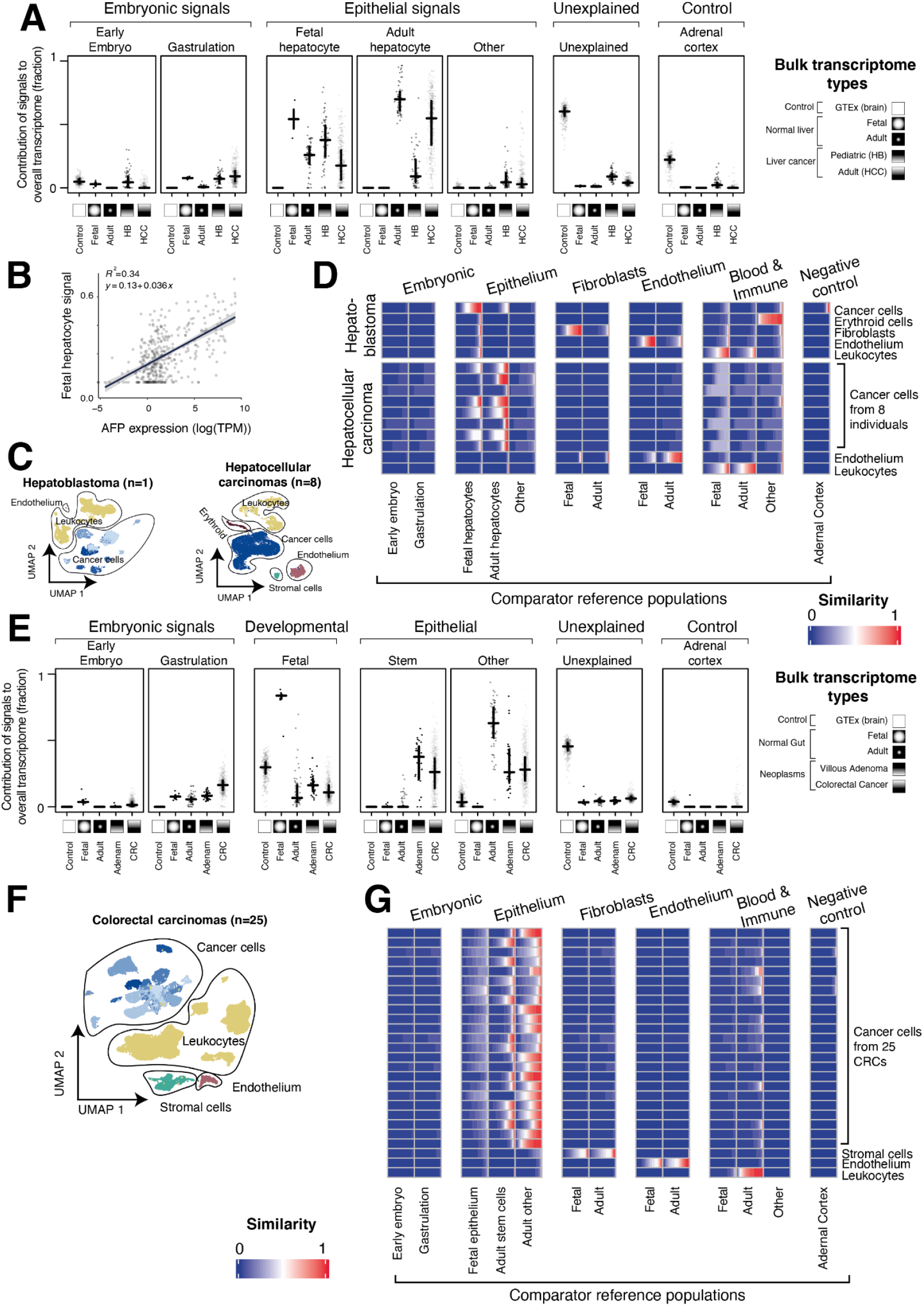
Detailed analysis of bulk and single cell liver and gut cancers. **A. Contribution of different signal types to bulk transcriptomes of liver:** Relative contribution (y-axis) of different reference single cell populations (horizontal facets) in explaining bulk transcriptomes (dots) grouped by transcriptome type (x-axis within facets) as indicated by x-axis symbols and labels. The fit was performed with all single cell reference atlases provided, related groups of reference cell populations indicated by hierarchical labels (top), and the distribution of groups of bulk transcriptomes summarised by their median (horizontal lines) and 1st/3rd quartiles (vertical lines). **B. Foetal hepatocyte contribution correlated with AFP expression:** Foetal hepatocyte contribution to bulk transcriptomes of hepatocellular carcinoma (y-axis) plotted against alpha fetoprotein (*AFP*) expression (log10 of TPM, x-axis), for each bulk transcriptome (dots). A best fit linear trend line is shown along with its equation and associated R squared value. The fit to the bulk transcriptome was performed excluding *AFP.* **C. UMAP of liver cancer transcriptomes:** Dimensionality reduction analysis (UMAP) showing single cell transcriptomes (dots) derived from 1 hepatoblastoma (left) and 8 individual hepatocellular carcinomas (right), grouped by cell type (contours, labels, and colours). Different donors of cancer cells are indicated by different shades of blue. **D. Similarity of single cell HCC/HB transcriptomes with embryonic and liver reference atlases:** Similarity score (logistic regression, colour value) of single cell transcriptomes of liver cancers (**C.**) grouped by cell type (y-axis) and compared to transcriptomes of reference cell types (x-axis) using a reference consisting of embryonic cells, developmental liver, and post-natal liver. Each rectangle represents a group of cells (indicated by y-axis label) and shows the distribution of similarity scores for those single cells compared to the reference cell population (indicated by x-axis label). **E. Contribution of different signal types to bulk transcriptomes of the intestines:** Relative contribution (y-axis) of different reference single cell populations (horizontal facets) in explaining bulk transcriptomes (dots) grouped by transcriptome type (x-axis within facets) as indicated by x-axis symbols and labels. The fit was performed with all single cell references provided, related groups of reference cell populations indicated by hierarchical labels (top), and the distribution of groups of bulk transcriptomes summarised by their median (horizontal lines) and 1st/3rd quartiles (vertical lines). **F. UMAP of colorectal carcinoma cell transcriptomes from 25 individuals:** Dimensionality reduction analysis (UMAP) showing single cell transcriptomes (dots) derived from 25 individual HCCs, grouped by cell type (contours, labels, and colours). Different donors of cancer cells are indicated by different shades of blue. **G. Similarity of single cell CRC transcriptomes with embryonic and intestine reference atlases:** Similarity score (logistic regression, colour value) of single cell transcriptomes of colorectal cancers (**F.**) grouped by cell type (y-axis) and compared to transcriptomes of reference cell types (x-axis) using a reference atlas consisting of embryonic cells, developmental intestine, and post-natal intestine. Each rectangle represents a group of cells (indicated by the label on the y-axis) and shows the distribution of similarity scores for those single cells compared to the reference cell population (indicated by x-axis label).

The next step in the analysis was to determine which precise liver cell type best explains the transcriptome in hepatoblastoma and hepatocellular carcinoma and to exclude that the signal derives from non-parenchymal cells. We found that adult and foetal hepatoblasts, alone or in combination, were the main cell signal in adult hepatocellular carcinoma (**Figure 2A, S5**). In some HCCs, the foetal hepatoblast signal predominated, which correlated with serum AFP levels and AFP mRNA counts. The predominance of this signal persisted when we removed AFP from the reference transcriptome, indicating that it was not driven by this transcript alone (**Figure 2B**). The foetal state of some hepatocellular carcinoma may be of prognostic significance, as AFP serum levels have previously been associated with poor outcomes in hepatocellular carcinoma^23^. Occasional adult tumours exhibited signals of other rare liver cells, namely foetal hepatobiliary hybrid progenitors or so called adult BEC (hepatocytes and biliary epithelial cells, **Figure S5**). A clinical or pathological significance of these unusual cellular signatures was not apparent. Consistent with our pan-tissue analysis (**Figure 1C**), we found that the foetal hepatocyte signal dominated in the childhood cancer hepatoblastoma (**Figure 2A**).

To validate these findings, we integrated (published) single cell transcriptomes from hepatocellular carcinomas^13^ (n = 25,605 cells from 8 tumours) and generated 13,180 single cell transcriptomes from a hepatoblastoma using the Chromium 10X platform (**Figure 2C**). For each single cell transcriptome we calculated a similarity score (using logistic regression^24^) against the same reference used to analyse the bulk transcriptomes (**Figure 2D**). The single cell mRNA analyses verified our bulk transcriptomic findings, namely that adult hepatocellular carcinoma may be viewed as aberrant adult hepatocytes, some of which dedifferentiate towards a foetal hepatoblast state (**Figure 2D**). By contrast, most childhood hepatoblastoma cells matched foetal hepatoblasts (**Figure 2D**).

We next applied our analytical approach to bulk and single cell transcriptomes of colorectal cancers, as well as to adenomas (polyps)^14,16^. Adenomas are low grade neoplastic lesions of colorectal epithelium that have the potential to progress to carcinomas, via the sequential acquisition of cancer-causing somatic mutations (adenoma-carcinoma sequence). In the first instance, we assessed which reference most fully accounted for adenoma and cancer transcriptomes (**Figure 2E, S6**) and again found that the tissue specific colorectal reference, but not more primordial references, provided the best fit. We then analysed which specific cell type of the colorectal tissue reference mostly explained adenoma bulk transcriptomes (**Figure 2E**). We found that adult, but not foetal, colorectal stem cell signals predominated across cancers, with a roughly comparable contribution from more mature epithelial cells. Interestingly, the same stem cell signal pervaded adenomas. Thus, transcriptional, genetic, histological differences notwithstanding, from a cell state perspective there was no obvious change (eg, further dedifferentiation) from adenomas to carcinomas. We next verified cell signals in colorectal cancers by comparing single cancer cells^14^ with normal reference cells, which confirmed that colorectal cancer cells resemble a mixture of stem and more mature epithelial cells (**Figure 2F,G**) that do not dedifferentiate to more primordial states. Overall these findings indicate that the predominant cell state of premalignant and malignant colorectal tumours is the colorectal stem cell. This may indicate that the cell of origin of these neoplasms is the colorectal stem cells or, that irrespective of where in the differentiation hierarchy of colorectal epithelium tumours originate, they ultimately converge at the stem cell state.

Our analyses reveal a nuanced picture of the differentiation state of adult cancers, wherein most adult tumours were best explained by post-natal cells, but not by a reversion to an antenatal state. By contrast, a “fetal-like” transcriptome was a near-universal feature of childhood tumours, likely as a consequence of their probable origins in development. These findings suggest that cancer dedifferentiation, at the level of the transcriptome, is a highly tissue and cancer type specific process, rather than a general hallmark of adult cancer.

## Methods

### Ethics statement

Informed consent for research was obtained from participants (or their carers). Studies underlying this paper have received appropriate approval by ethics review boards as per national legislation. U.K. tumour samples were collected under the following study: National Health Service (NHS) National Research Ethics Service reference 16/EE/0394.

### Single-cell hepatoblastoma processing

Surplus tumour tissue obtained at diagnostic biopsy or tumour resection was processed immediately after receipt in the histopathology laboratory (<1 hour after interventional radiology/surgical procedure). Tissue was minced using a scalpel and then incubated in RPMI 1640, supplemented with 10% fetal calf serum, 1% l-glutamine, and 1% peni- cillin/streptomycin, with collagenase IV (1.6 mg/ml; catalog no. 11410982; MP Biomedi- cals), for 30 min at 37°C, inverting the tube every 10 min. The digested tissue was passed through a 70-μm filter and incubated in 1 x RBC lysis buffer (catalog no. 420301; BioLe- gend) for 10 min at room temperature.

The obtained single-cell suspension was used for downstream processing. Part of the single-cell suspension was depleted of CD45+ cells to enrich for tumour cells using a CD45 MicroBeads kit (catalog no. 130-045-801; Miltenyi Biotec), following the manufacturer’s protocol. Both CD45 nondepleted and CD45-depleted single-cell suspensions were depleted of dead cells using a Dead Cell Removal kit (catalog no. 130-090-101; Miltenyi Biotec), following the manufacturer’s protocol. Obtained viable single-cell suspensions were processed on the 10x Chromium platform.

The concentration of single-cell suspensions was manually counted using a haemocy- tometer and adjusted to 1000 cells/μl or counted by flow cytometry. Cells were loaded according to the standard protocol of the Chromium Single Cell 3’ Kit (v2 and v3 chem- istry). All the following steps were performed according to the standard manufacturer’s protocol. One lane of Illumina HiSeq 4000 per 10x chip position was used.

Single-cell RNA-seq data were mapped, and counts of molecules per barcode were quanti- fied using the 10x software package cellranger (versions 2.0.2 and 3.0.2) to map sequencing data to version 2.1.0 of the build of the GRCh38 reference genome supplied by 10x.

### Single-cell data processing

To perform cell signal analysis and/or logistic regression, the following were required for each single-cell dataset: i) a count table that has undergone quality control; ii) cell annotations. Where both could be obtained, no further action was taken. These were taken directly from the publication where available, or reproduced following the methods of the relevant publication. All datasets were 10x, except where specified below. All cell types with <10 cells were removed. When merging single cell data matrices from different sources, only common rows (ENSEMBL IDs wherever possible) were kept. Additional processing was performed for the following references:

#### Gastrulation data

QC’d count tables and annotations were obtained from the authors. As this was a Smart-Seq2 dataset, the raw count table was transformed to transcripts per kilobase million (TPM) to approximate UMI values of 10x datasets.

#### Fetal liver

QC’d count tables and annotations were obtained from the authors. Sub-types of erythroid cells, B-cells and dendritic cells were merged into one cell type to simplify the annotation (e.g. early, mid, and late erythroid cells were re-labelled to erythroid cells).

#### Adult liver

QC’d count tables and annotations were obtained from the authors. Clusters of hepatocytes, T-cells and liver sinusoidal endothelial cells into one cell type to simplify the annotation.

#### Epithelial liver cell reference by Segal et al

A QC’d raw count table was obtained from the authors. As this was a Smart-Seq2 dataset, raw count tables were transformed to transcripts per kilobase million to approximate UMI values of 10x datasets. Adult and fetal HHyP and hepatocyte, and adult BECs, were annotated based on marker gene expression.

#### Hepatoblastoma

Single-cell RNA-seq data were mapped, and counts of molecules per barcode were quantified using the 10x software package cellranger (version 3.0.0) to map sequencing data to version 2.1.0 of the build of the GRCh38 reference genome supplied by 10x. Cells with >20% mitochondrial expression, fewer than 200 detected genes, or 500 UMIs were removed as low quality. Data were log normalised and clustered using a community detection method^25^. Clusters were annotated with the following markers: *CD45* (leukocyte marker), *HBA* and *HBB* (erythrocyte markers), *EPCAM* and *AFP* (tumour cell markers),*PECAM1* (endothelial marker) and *ACTA2* (hepatic stellate marker). Only those cells that could be definitively annotated using these markers were used for the analysis.

#### Hepatocellular carcinoma

Single-cell RNA-seq data were mapped, and counts of molecules per barcode were quantified using the 10x software package cellranger (version 3.0.0) to map sequencing data to version 2.1.0 of the build of the GRCh38 reference genome supplied by 10x. Cells with >10% mitochondrial expression, fewer than 300 detected genes, or 1000 UMIs were removed as low quality. Cells were then annotated using marker gene expression^13^.

### Bulk data processing

To perform cell signal analysis, the following were required for each bulk sample i) gene counts; ii) gene lengths. These were generated in the following manner:

#### TCGA data

Gene counts and lengths were taken from the recount2 mapping of the TCGA^26^.

#### St Jude’s

Raw sequencing reads were mapped against the GRCh38 reference v1.2.0 provided by 10X, using a pseudo-aligner^27^, which produced both gene counts and lengths. Cancers which were not unique to childhood (osteosarcoma and melanoma) were removed.

#### Hepatoblastoma (tumour and normal)

Gene counts were provided by authors, the same gene lengths as used for the TCGA were used.

#### Colorectal bulk (tumour, normal, adenomas)

Gene counts and lengths were provided by the authors.

#### Fetal liver bulk

Gene counts were provided by authors, the same gene lengths as used for the TCGA were used.

#### Fetal gut bulk

Gene counts were provided by authors, the same gene lengths as used for the TCGA were used.

#### Blastoid bulk

Gene counts were downloaded from GEO, at accession GSE179040, the same gene lengths as used by the TCGA were used.

#### GTEx

TPM values provided by the GTEx consortium were used, with gene lengths set to 1.

### Cell signal analysis: comparing bulk transcriptomes to a single cell reference

Cell signal analysis was performed as previously described^18^. For the pan-cancer analysis, we excluded mitochondrial and ribosomal genes, and downweighted the likelihood for housekeeping genes by 50%. From the fetal reference, we excluded reference cell types from the fetal heart as there were no corresponding cell types in the adult reference.

### Cell similarity: comparing single cell transcriptomes to a single cell reference

To measure the similarity of a target single-cell transcriptome to a reference single-cell dataset, logistic regression was used to train a model on the reference single cell dataset^24^. This model was then applied to predict the probability of similarity (normalised to 1 across all categories), between each cell type in the reference dataset and each cell in the target dataset.

### Stemness score calculation

Code to calculate the stemness score as previously described was provided by the authors^4^. To provide a comparison between all samples, both adult and paediatric, the stemness score was calculated across all bulk samples from both the TCGA and St Judes simultaneously.

### Cell signal summary score calculation

Cell signal analysis produces for each bulk transcriptome, the relative contribution of each single cell defined cell type provided in the reference. In this paper, we make use of various summary scores (embryoness, fetalness, etc) which we define as the sum of all relevant cell signal analysis scores for a bulk transcriptome. For example, we defined the fetalness score as the sum of all the cell signal contributions across all cell types derived from fetal tissues, when cell signal analysis is performed using a reference consisting of both a fetal and mature tissue reference.

## Data availability statement

The data used in this study was obtained from the public sources indicated in **Table S1-2** and processed as described in the methods section. Additionally, single cell hepatoblastoma transcriptomes were generated and can be obtained from the EGA under accession number XXX.

## Code availability statement

The code used to process the data and generate these figures has been made available in a public repository, accessible here XXXX.

## Author contribution

M.D.Y., S.B., and I.C-C conceived and supervised the project, and wrote the manuscript. G.K., M.K., S.v.D., and M.D.Y. performed the analysis and data processing. C.T., A.P., K.A., E.P., and K.S. collected samples and generated data. S.T., M.H., and R.A. contributed to understanding the reference data.

## Supplementary Figures

**Supplementary Figure 1.**
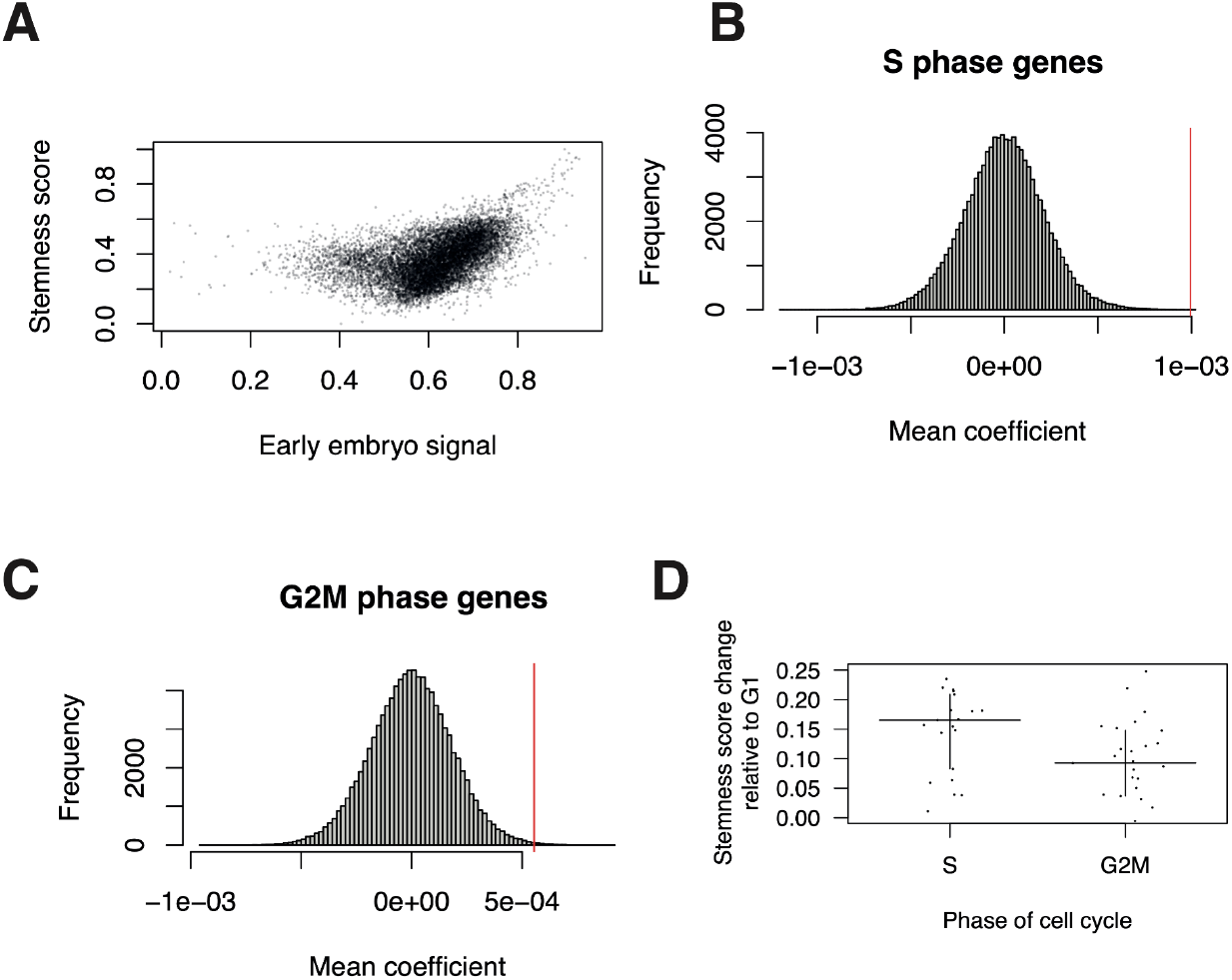
Relationship between stemness and early embryo scores. A. **Correlation between stemness and early embryo scores:** Stemness score (y-axis) as a function of early embryo signal (x-axis), for bulk cancer transcriptomes (dots). **B. Significance of S phase genes in stemness score:** The distribution across many randomly selected gene sets (y-axis) of average coefficients for that gene set in determining the stemness score (x-axis). Values further away from zero mean that a set of genes has a consistent contribution to determining the stemness value. The value corresponding to the set of genes linked to S-phase of the cell cycle is marked with a red line. **C. Significance of G2M phase genes in stemness score:** The same as **B.** but using genes associated with G2 and M phase of the cell cycle. **D. Shift in stemness score with cell cycle phase:** The single cell fetal liver reference data was used to construct pseudo-bulk counts from which stemness scores were calculated. For each cell type, a separate pseudo-bulk was constructed for cells in each phase of the cell cycle. The difference in stemness score for each cell type (y-axis) was calculated between other phases of the cell cycle (x-axis) and G1 phase. Horizontal lines indicate the median shift across all cell types, while the vertical lines indicate the 1st and 3rd quartiles.

**Supplementary Figure 2.**
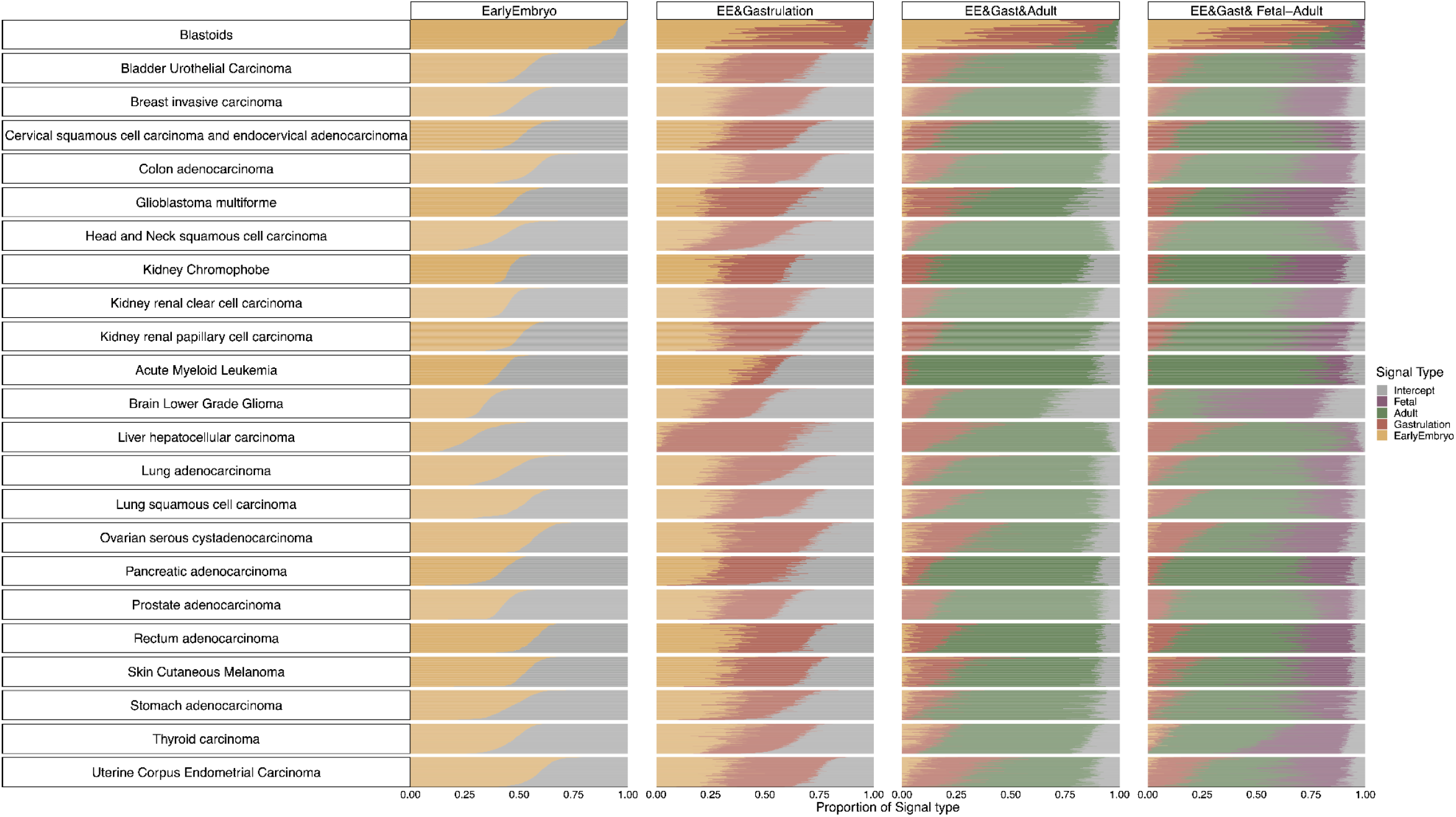
Cell signal analysis of all TCGA samples. Relative contribution of single cell reference populations (x-axis) in explaining all TCGA bulk adult cancer transcriptomes (y-axis) using cell signal analysis for different combinations of reference populations (columns splits, with heading indicating reference combination). Individual cancer transcriptomes (rows) are split into groups based on cancer types, as indicated by the row split titles. Colours indicate the contribution to each transcriptome of each reference type, as described by the legend on the right.

**Supplementary Figure 3.**
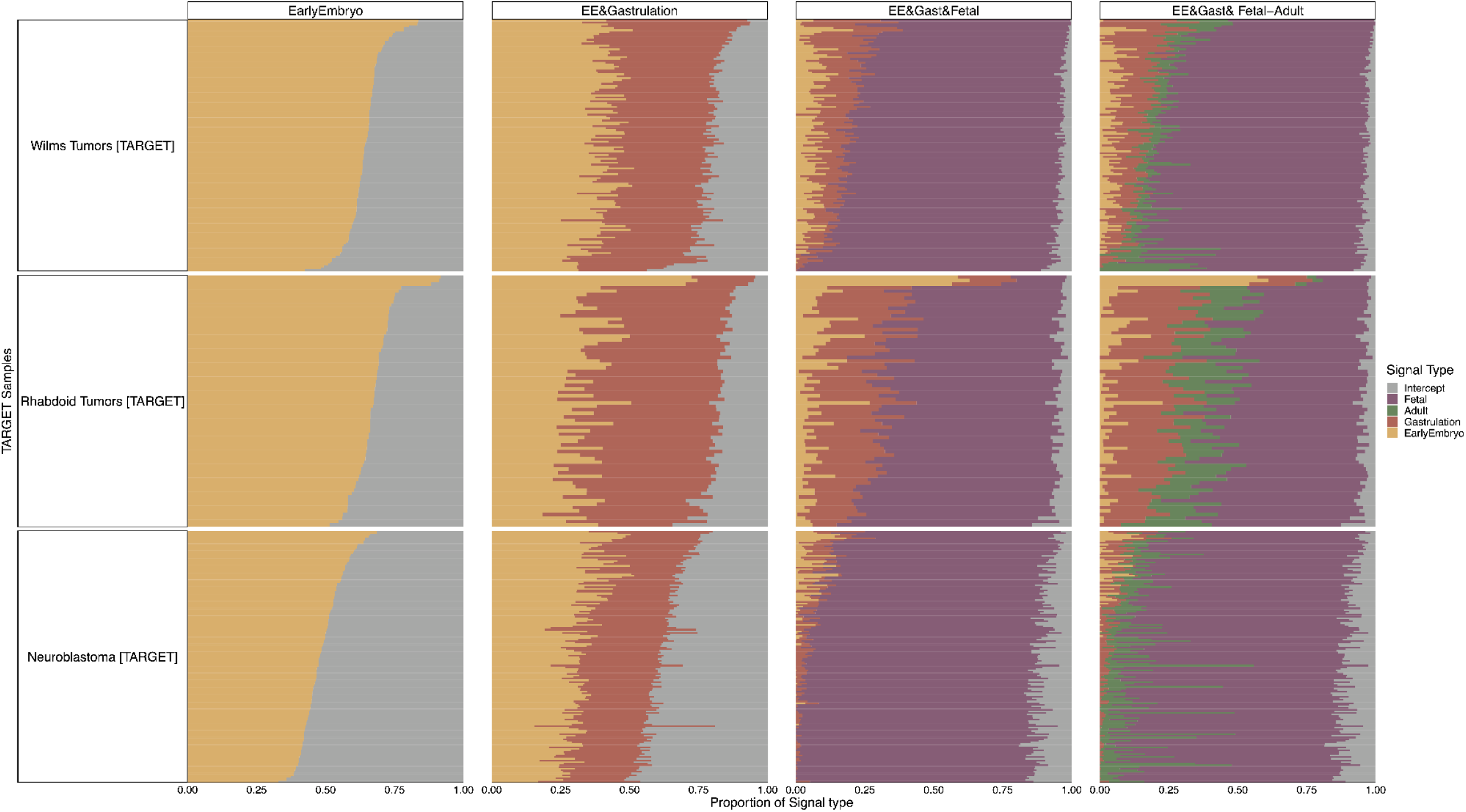
Cell signal analysis of all TARGET samples. Relative contribution of single cell reference populations (x-axis) in explaining all TARGET bulk childhood cancer transcriptomes (y-axis) using cell signal analysis for different combinations of reference populations (columns splits, with heading indicating reference combination). Individual cancer transcriptomes (rows) are split into groups based on cancer types, as indicated by the row split titles. Colours indicate the contribution to each transcriptome of each reference type, as described by the legend on the right.

**Supplementary Figure 4.**
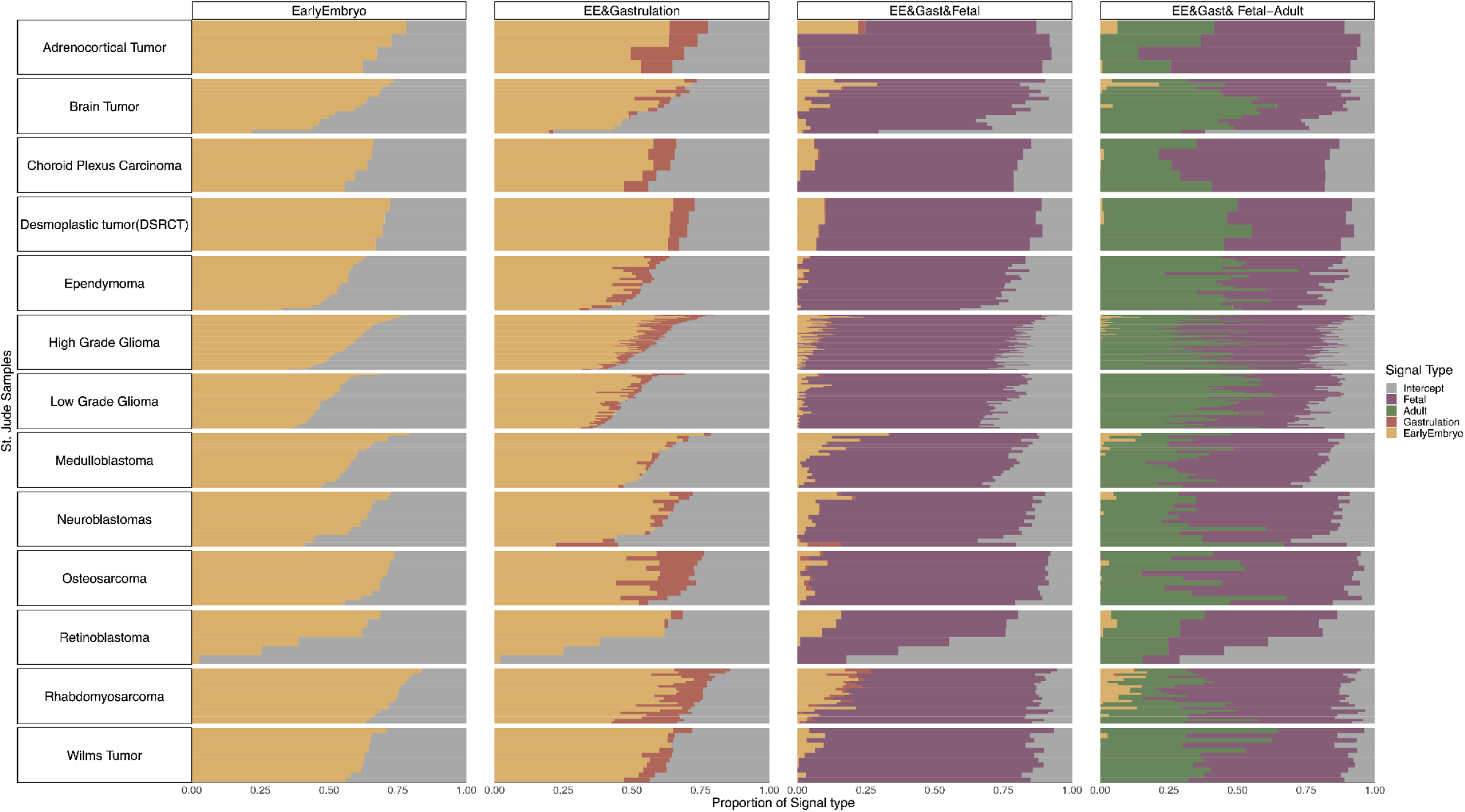
Cell signal analysis of all StJudes samples. Relative contribution of single cell reference populations (x-axis) in explaining all StJudes bulk childhood cancer transcriptomes (y-axis) using cell signal analysis for different combinations of reference populations (columns splits, with heading indicating reference combination). Individual cancer transcriptomes (rows) are split into groups based on cancer types, as indicated by the row split titles. Colours indicate the contribution to each transcriptome of each reference type, as described by the legend on the right.

**Supplementary Figure 5.**
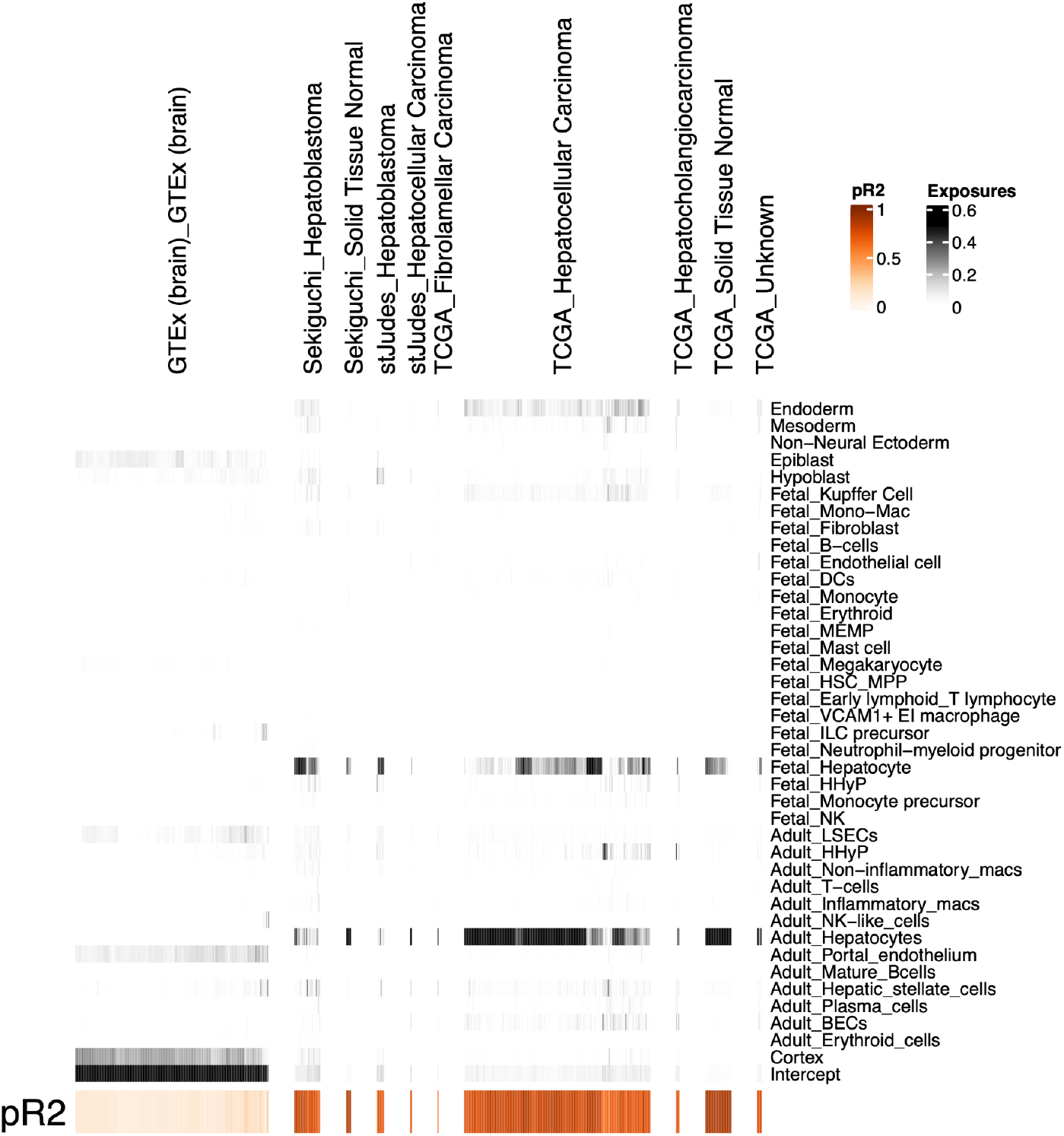
Cell signal analysis of all liver samples. Relative contribution of single cell populations from the fetal and adult liver, gastrulation, and early embryo (y-axis) in explaining bulk transcriptomes (y-axis) using cell signal analysis. Bulk transcriptomes (columns) are grouped together into the categories indicated by the top labels and the relative contribution in explaining the signal is indicated by the intensity of the greyscale as indicated by the legend. For each bulk transcriptome, a goodness of fit (pseudo R squared) was calculated and is indicated by the intensity of the orange colour in the bottom row, as described by the legend.

**Supplementary Figure 6.**
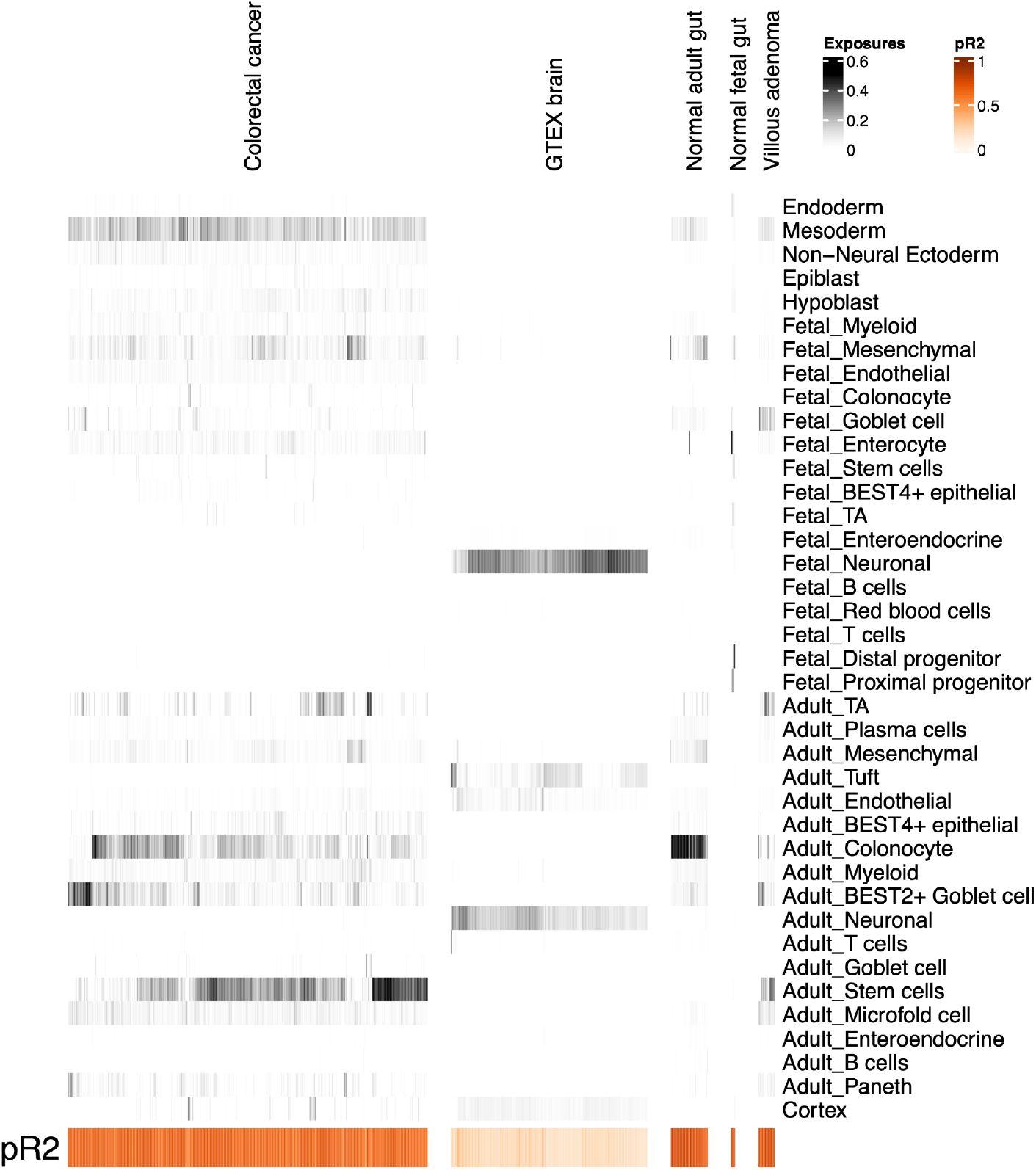
Cell signal analysis of all gut samples. Relative contribution of single cell populations from the fetal and adult gut, gastrulation, and early embryo (y-axis) in explaining bulk transcriptomes (y-axis) using cell signal analysis. Bulk transcriptomes (columns) are grouped together into the categories indicated by the top labels and the relative contribution in explaining the signal is indicated by the intensity of the greyscale as indicated by the legend. For each bulk transcriptome, a goodness of fit (pseudo R squared) was calculated and is indicated by the intensity of the orange colour in the bottom row, as described by the legend.

## Supplementary Tables

**Supplementary Table 1**

Table summarising all single cell transcriptomic datasets used in this study, along with the source from which they were obtained.

**Supplementary Table 2**

Table summarising all bulk transcriptomic samples used in this study, along with the source from which they were obtained.

